# Time-resolved proximity proteomics uncovers a membrane tension-sensitive caveolin-1 interactome at the rear of migrating cells

**DOI:** 10.1101/2022.12.13.520222

**Authors:** Rossana Girardello, Eleanor Martin, Gunnar Dittmar, Alexander Ludwig

## Abstract

Caveolae play fundamental roles in mechanotransduction. Critical to caveolae function is their ability to flatten out in response to an increase in membrane tension, thereby acting as a membrane reservoir to buffer acute mechanical stress. Cycles of caveolae assembly and disassembly also regulate membrane tension at the rear of migrating cells via RhoA/ROCK-mediated actomyosin contractility. However, the molecular mechanisms that couple caveolae-mediated mechanotransduction to cortical actin dynamics are poorly understood. Here we used biotin-based proximity labelling and quantitative mass spectrometry to define a caveolae-associated interactome in migrating RPE1 cells at steady state and in response to an acute increase in membrane tension induced by hypo-osmotic shock. Our data reveal a dynamic caveolae-associated protein network composed of focal adhesion proteins and cortical actin regulators that is highly sensitive to changes in membrane tension. We show that membrane tension differentially controls the association of ROCK and the RhoGAP ARHGAP29 with caveolae and that ARHGAP29 regulates caveolin-1 Y14 phosphorylation, caveolae rear localisation and RPE1 cell migration. Caveolae in turn regulate ARHGAP29 expression, most likely through the control of YAP signalling. Taken together, our work uncovers a membrane tension-dependent functional coupling between caveolae and the rear-localised actin cytoskeleton, which provides a framework for dissecting the molecular mechanisms underlying caveolae-regulated mechanotransduction pathways.

## Introduction

Caveolae are an abundant feature of the plasma membrane of almost all vertebrate cells. They are composed of the scaffolding proteins caveolin-1 (Cav1) and caveolin-2 (Cav2), which are embedded in the inner leaflet of the plasma membrane, and the cavin protein family, which form a loose peripheral caveolar membrane coat. There are four cavins in humans (cavin1/PTRF, cavin2/SDPR, cavin3/SRBC and cavin4/MURC). Cav1 and cavin1 are essential for caveolae formation, whereas cavins 2-4 regulate caveolae dynamics and functions in a cell and tissue-specific fashion (1–4). A third caveolin protein (Cav3) and cavin4 are expressed exclusively in muscle cells. In addition, the ATPase EHD2 and the F-BAR domain containing protein Pacsin2 are found at caveolae. While caveolins and cavins assemble into an oligomeric membrane coat that shapes the caveolar bulb (3, 5–9), EHD2 and Pacsin2 are localised to the caveolar neck and link caveolae to actin filaments to restrict their mobility (8, 10–13).

Caveolae are extraordinarily dynamic in nature and have been implicated in diverse biological processes (14, 15). Caveolae are abundant in cells that are exposed to tensile forces or shear-stress, such as endothelial cells, epithelial cells, and muscle cells, and protect such cells from mechanical stress (16–22). There is evidence that caveolae disassemble (or flatten out) in response to an increase in membrane tension (23), and may similarly respond to other stresses such as osmotic stress (24), oxidative stress (25), and UV irradiation (26). Tension-induced flattening was proposed to be mediated by the dissociation of the peripheral cavin coat and EHD2 from membrane-embedded caveolin oligomers (23, 27). Cavin complexes and EHD2 released into the cytosol can subsequently translocate into the nucleus to regulate gene transcription (26, 28, 29). This suggests that caveolae act as a dynamic mechanosensitive membrane reservoir that senses changes in membrane tension and transmits such inputs to downstream signaling (30–34). However, temporal and molecular insight into how caveolae respond to membrane tension is limited.

Large numbers of caveolae are also found at the rear of migrating cells, which exhibit an intrinsic front-rear gradient in membrane tension (35, 36). Several studies indicate an intricate relationship between caveolae-mediated mechanosensing and RhoA-mediated cell rear retraction (37–39). Caveolae formation at the cell rear is promoted by low membrane tension and is dependent upon RhoA/ROCK1 signaling, the Rho guanidine nucleotide exchange factor (GEF) Ect2, and the RhoA effector protein and serine-threonine kinase PKN2. Caveolae formation, in turn, enhances RhoA/ROCK1 signaling, leading to F-actin alignment, actomyosin contractility and rapid rear retraction (39, 40). Loss of caveolae, increased membrane tension, or inhibition of Rho signalling all break this positive feedback loop, inhibiting rear retraction and impeding cell migration (39, 40). This indicates a central role for caveolae in coupling changes in membrane tension to the control of actomyosin contractility at the rear of migrating cells. However, the molecular machinery that regulates the assembly and disassembly of caveolae at the cell rear is not well defined.

Here we employed proximity biotinylation with the peroxidase APEX2 and quantitative proteomics to rapidly label and measure the Cav1-associated protein network in migrating RPE1 cells at iso-osmotic conditions and in response to an acute increase in membrane tension elicited by hypo-osmotic shock. We define a caveolae-associated protein network at the rear of migrating cells that is highly sensitive to changes in membrane tension. We also identify a number of potentially novel regulators and effectors of caveolae and show that one of them, the Rho GTPase activating protein (GAP) ARHGAP29, controls caveolae rear localisation and RPE1 cell polarity and migration.

## Results and Discussion

### Caveolae are abundant at and stably associated with the rear of RPE1 cells

We previously demonstrated that caveolae are enriched at the rear of hTERT-RPE1 cells (8). To study the dynamics of caveolae in migrating RPE1 cells we used a cell line stably transfected with a cavin3-miniSOG-mCherry fusion protein (8). Time lapse microscopy of cells migrating on a fibronectin-coated glass surface showed that caveolae were stably associated with the cell rear for several hours (Figure 1A and Supplemental Movie S1). Instantaneous rear retractions were observed. Moreover, caveolae were stably associated with the rear of RPE1 cells migrating in a 3D collagen matrix (Figure 1B and Supplemental Movie S2), as shown previously in A2780 cells (39). Next we used miniSOG labeling (8, 41) and electron tomography to visualise the 3D architecture of caveolae. 3D surface rendering revealed a dense network of surface-connected caveolae and the presence of large interconnected caveolar clusters, most of which appeared to be continuous with the rear plasma membrane (Figure 1C and Supplemental Movie S3). We concluded that caveolae are abundant at the rear of RPE1 cells, which hence provide an appropriate model cell type to study the caveolae-associated protein network in migrating cells under normal conditions and in cells experiencing high membrane tension.

**Figure 1.**
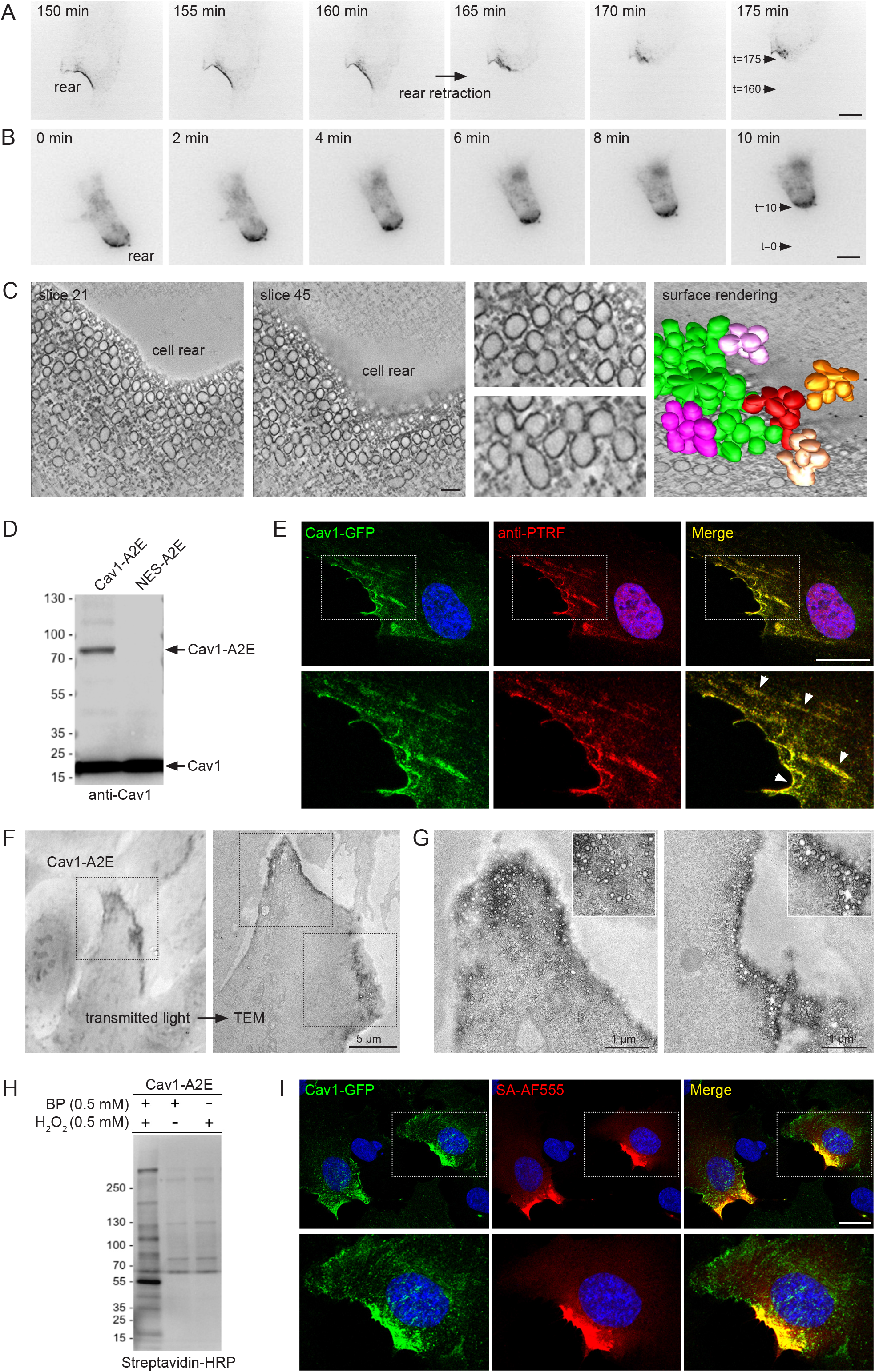
Dynamics and 3D architecture of caveolae at the rear of migrating RPE1 cells. A) Time lapse images of RPE1 cells stably transfected with cavin3-miniSOG-mCherry migrating on a glass coverslip. A) Time lapse images of RPE1 cells stably transfected with cavin3-miniSOG-mCherry migrating on in a collagen matrix. C) Electron tomography and surface rendering of caveolar networks at the rear of RPE1 cells stably transfected with cavin3-miniSOG-mCherry. D) Western blot of stable RPE1 cell lines expressing either Cav1-APEX2-EGFP or NES-APEX2-EGFP fusion proteins. The membrane was probed with anti-Cav1 antibodies. E) Confocal fluorescence microscopy of Cav1-A2E RPE1 cells stained with anti-PTRF/cavin1 antibodies. Boxed regions are magnified in the bottom panel. Nuclei were counterstained with DAPI. Scale bar 10 μm. F) Transmitted light image of Cav1-A2E RPE1 cells after the APEX2 labeling reaction and plastic embedding (left panel) and correlative TEM image of the same cell (right panel). G) Representative TEM micrographs of serial TEM sections of the cell shown in C (boxed regions). Note the abundance of caveolae at the cell rear. H) Cav1-APEX2-EGFP expressing cells were incubated in the absence or presence of biotin phenol and/or hydrogen peroxide. Cell lysates were separated by SDS-PAGE, blotted onto PVDF membrane, and probed with streptavidin-HRP. I) Confocal fluorescence microscopy of Cav1-APEX2-EGFP RPE1 cells after proximity labelling with APEX2. Cells were fixed and stained with fluorescent streptavidin (Alexa Fluor 555) to visualize biotinylated proteins. Nuclei were counterstained with DAPI. Scale bar 10 μm.

### The caveolin-1 interactome is enriched in cortical actin regulators and is regulated by membrane tension

Several cytoplasmic effectors of cavins have been identified using BioID-mediated proximity proteomics (26, 29, 42). However, BioID requires long labeling times (16-24 hours) and therefore is not suitable to dissect rapid changes in the caveolae-associated protein network that are likely to occur as cells respond to acute mechanical stress. To address this we used proximity biotinylation with the peroxidase APEX2 (43, 44). A stable RPE1 cell line expressing a Cav1-APEX2-EGFP (Cav1-A2E) fusion protein was generated. We have previously shown that expression of this construct rescues caveolae formation in mouse embryonic fibroblasts from *CAV1* -/- mice, indicating that this fusion protein is functional (7, 45). Cav1-A2E in RPE1 cells was expressed at levels comparable to or lower than that of endogenous Cav1 (Figure 1D) and colocalised with PTRF/cavin1 at the cell rear, as expected (Figure 1E). Moreover, transmission electron microscopy showed that the Cav1-A2E fusion protein was efficiently incorporated into caveolae (Figure 1F and 1G). Next we ascertained that proximity biotinylation in the Cav1-A2E cell line is specific and spatially restricted. Streptavidin-HRP blotting of Cav1-A2E RPE1 cell lysates showed that the biotinylation reaction was dependent upon the presence of biotin phenol and a brief (1 min) exposure to hydrogen peroxide (Figure 1H). In addition, fluorescent streptavidin labeling showed that the biotinylation reaction was restricted to the cell rear and colocalised with Cav1-A2E (Figure 1I). We also generated a control RPE1 cell line expressing APEX2-EGFP fused to a nuclear export signal (NES-A2E). In this cell line biotinylated proteins were diffusely localised in the cytoplasm, as shown previously (Figure S1) (46).

To generate Cav1 proximity proteomes at resting conditions and upon an acute increase in membrane tension we used a hypo-osmotic shock assay (23) (Figure 2A). Cav1-A2E expressing cells were either grown in isotonic medium and left untreated (non-treated; NT), exposed to a brief 5 min hypo-osmotic shock (HYPO), or exposed to hypo-osmotic shock and allowed to recover in iso-osmotic medium for 30 min (REC). NES-A2E cells grown in iso-osmotic medium (NT) were used as a reference (or “spatial ruler”) to subtract abundant cytoplasmic proteins and non-specific bystanders. After ratiometric filtering over the NES sample we identified a total of 348 unique proteins that were significantly enriched in the three Cav1 proximity proteomes. 70 proteins were exclusive to the HYPO sample, 15 proteins were enriched specifically in the HYPO and REC samples, and 26 proteins were enriched specifically in the NT and REC samples (Figure 2B and Figure S2). Importantly, Cav1-A2E and all major caveolar proteins (Cav1, Cav2, PTRF/cavin1 and EHD2; note that cavin2 and cavin3 are not expressed in RPE1 cells) were specifically enriched in the iso-osmotic control (NT) and recovered (REC) samples. On the contrary, under hypo-osmotic (HYPO) conditions Cav1, Cav2, and EHD2 were no longer significantly enriched with Cav1-A2E (Figure 2C), suggesting that high membrane tension had caused the dissociation of these proteins from Cav1-A2E. Indeed, Cav1-A2E, PTRF/cavin1, and EHD2 were all specifically enriched in the NT and REC samples when directly compared to the HYPO sample (Figure 2D). By contrast, no significant differences in the abundance of caveolar core components were observed between NT and REC samples, suggesting that caveolae formation is fully restored within 30 min of recovery from a hypo-osmotic shock. We concluded that proteins identified specifically in the NT and REC samples are associated with Cav1 in a membrane-tension-dependent manner and therefore constitute a reliable Cav1 proximity proteome under iso-osmotic conditions.

**Figure 2:**
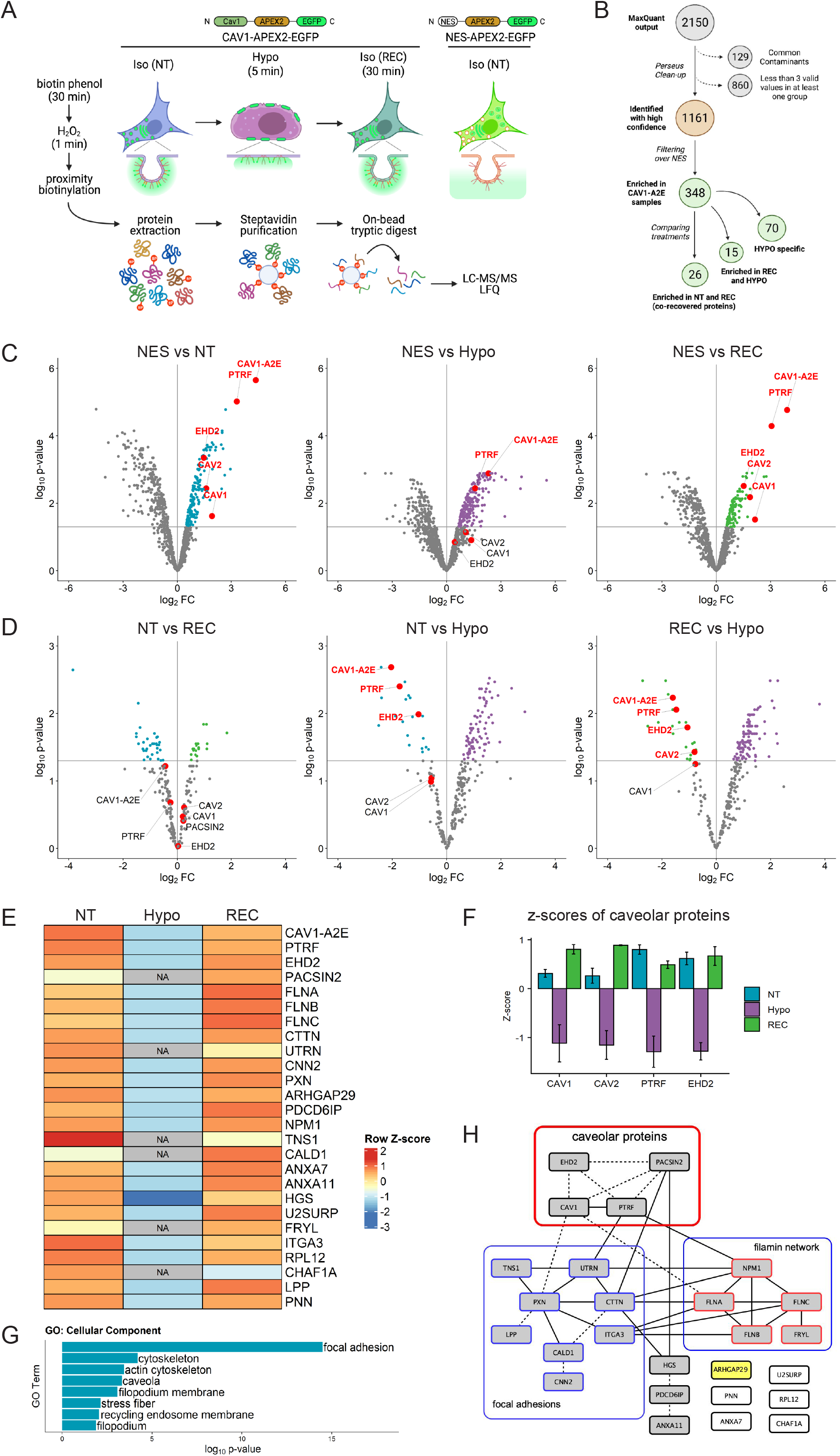
The caveolin-1 interactome is enriched in cortical actin regulators and focal adhesion proteins and is highly sensitive to membrane tension. A) Experimental set-up used for APEX2-mediated proximity biotinylation and LC-MS/MS. After proximity labelling, cells were lysed and biotinylated proteins were captured using magnetic streptavidin beads. Purified proteins were on bead-digested with trypsin and the recovered peptides analysed by LC-MS/MS. Three individual replicates were analysed. B) Diagram summarizing the LC-MS/MS analysis and the total number of proteins significantly enriched in each treatment compared to the NES control. Only proteins identified in all three replicates in at least one of the three samples were considered for further analysis. Label-Free Quantification (LFQ) was used to define specific Cav1 interactomes and to determine quantitative changes upon hypo-osmotic shock and recovery. C) Volcano plots showing proteins significantly enriched in NT, HYPO and REC samples compared to the NES control sample. Only proteins with a p-value <0.05 and a logFC>0 were considered for further analysis. D) Volcano plots comparing the NT and REC, NT and HYPO, and REC and HYPO samples. Cav1-A2E fusion protein and the core caveolar proteins are highlighted. E) Heatmap of z-scored LFQ quantified protein groups differentially enriched across the treatments. NA indicates not identified. See Figure S2 for a complete analysis. F) Z-scores of caveolar proteins under NT, HYPO and REC conditions. G) Gene Ontology (GO) Cellular Component analysis for proteins enriched under NT and REC conditions (see E). H) PPI network of the Cav1 proximity proteome. Only proteins significantly enriched in both the NT and REC samples were considered (i.e. 26 co-recovered proteins). Solid lines indicate interactions retrieved from databases (Biogrid, Reactome, BioCarta), dotted lines indicate interactions retrieved from a hand-curated literature search (see Table S1).

Quantitation of the MS data and GO annotation analyses revealed that the Cav1 proximity proteome was enriched in components of the cortical actin cytoskeleton and integrin adhesions, including filamins (FLNA, FLNB, FLNC), utrophin (UTRN), tensin-1 (TNS1), cortactin (CTTN) and paxillin (PXN) (Figure 2E–2G, Figure S2). A comprehensive database and literature review further identified several direct and indirect interactions between caveolar proteins, filamins and focal adhesion components (Figure 2H). These data support previous work demonstrating a role for caveolae in focal adhesion turnover and integrin-mediated mechanotransduction (47–50). Moreover, the F-actin crosslinking protein filamin-A and utrophin are well-known mechanotransducers (51, 52) and have been shown to regulate caveolae trafficking and dynamics, possibly by linking caveolae to the actin cytoskeleton via direct or indirect interactions with Cav1 or cavin1 (53–57). Hence, time-resolved proximity labelling with APEX2 uncovered a dynamic rear-localised Cav1 interactome enriched in mechanosensitive cortical actin regulators that is highly sensitive to acute changes in membrane tension.

To validate our proteomics data we quantified the spatial proximity between Cav1 and some of the identified proteins using proximity ligation assays (PLA) (Figures 3A–3F and Figure S3). In agreement with our proteomics data, under iso-osmotic conditions we observed robust PLA signals for Cav1-A2E and cavin1, Cav1-A2E and filamin-A, and Cav1-A2E and cortactin (Figure 3A–3D). Moreover, robust signals were observed for endogenous Cav1 and an APEX2-EGFP fusion protein of EHD2 (EHD2-A2E) (Figures 3E and 3F). PLA signals between Cav1-A2E and cavin1 and anti-Cav1 and EHD2-A2E were largely confined to the cell rear, as expected. Importantly, hypo-osmotic shock drastically reduced the PLA signals in all of the above cases, whereas cells that had recovered from the hypo-osmotic shock showed PLA signals indistinguishable to that of control cells. Taken together our data indicate that a brief hypo-osmotic shock causes an instantaneous but reversible disassembly of caveolae, as shown previously (23, 27, 28). The data further imply that membrane tension regulates the linkage between caveolae and the cortical actin cytoskeleton and that caveolae assembly/disassembly is spatio-temporally coordinated with the reorganisation of the rear-localised actin cortex.

**Figure 3:**
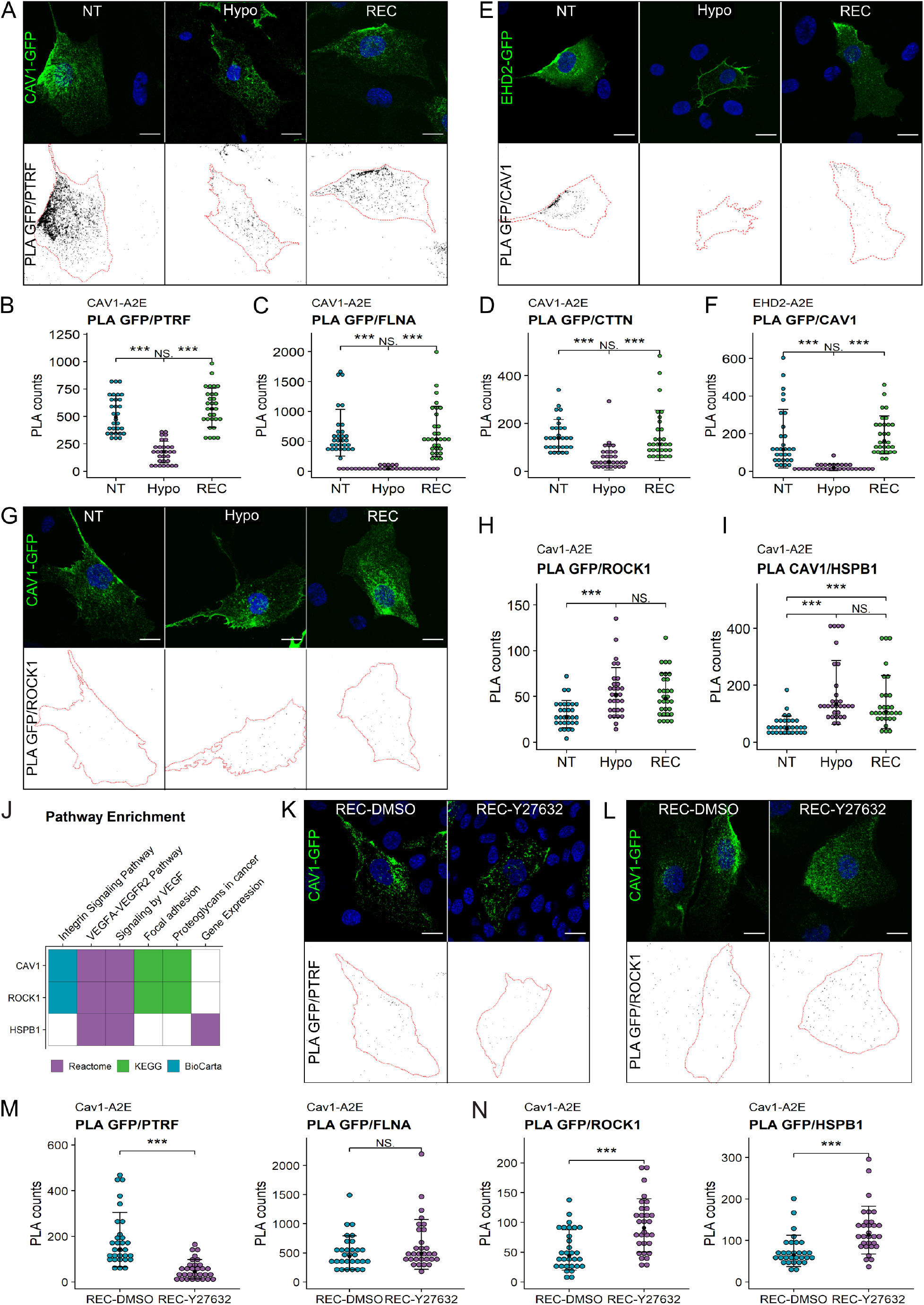
Proximity ligation assays reveal dynamic interactions between caveolae and the cortical actin cytoskeleton during changes in membrane tension. A-D) Proximity Ligation Assays in the Cav1-A2E RPE1 cell line using anti-GFP and anti-PTRF/cavin1 (A,B), anti-FLNA (C) or anti-CTTN (D) antibodies as indicated. E-F) Proximity Ligation Assays in RPE1 cells transfected with EHD2-A2E using anti-GFP and anti-Cav1 antibodies. G-I) Proximity Ligation Assays in the Cav1-A2E RPE1 cell line using anti-GFP and anti-ROCK1 (H) or anti-HSPB1 antibodies (I). J) Pathway enrichment analysis of proteins that are significantly enriched in the REC and HYPO samples compared to NT (See Heatmap in Figure 2E and Figure S2). K-N) Effect of ROCK inhibition on Proximity Ligation Assays in the Cav1-A2E RPE1 cell line using the indicated combinations of antibodies. Cells were incubated for 5 min in hypo-osmotic medium containing 10 μM Y-27632 or DMSO (control) and then transferred to isotonic medium for 30 min in the absence (DMSO) or presence of Y-27632. B,C,D,F,H,I,M,N) Quantifications of the PLA signals per cell. 30 cells from three independent experiments were quantified for each treatment. The number of PLA dots per cell is presented with the mean values ± SD. The bottom panels in A, E, G, K and L show the PLA signal masks used for counting. All scale bars are 20 μm.

### ROCK activity is required for caveolae reassembly upon hypo-osmotic shock

Interestingly, several proteins were specifically enriched with Cav1 during the recovery phase (i.e. in HYPO and REC samples) (Figure S2). Among these proteins were Rho-associated protein kinase 1 (ROCK1) and the heat shock protein HSPB1. Rho/ROCK1 signaling was previously shown to be required for caveolae localisation to the rear of migrating cells (39) and HSPB1 is a stress-regulated actin binding protein implicated in mechanotransduction (58, 59). Pathway enrichment analysis suggested a functional link between Cav1, ROCK1, and HSPB1 (Figure 3J). PLA assays confirmed that both HSPB1 and ROCK1 are significantly enriched with caveolae in cells exposed to hypo-osmotic shock and in cells that had recovered from the shock (Figures 3H–3I). To test whether ROCK1 activity, and hence actomyosin contractility, is required for caveolae reformation following hypo-osmotic shock, we performed PLA assays in the absence or presence of the ROCK inhibitor Y27632. ROCK inhibition blocked caveolae rear localisation (Figures 3K and 3L) and significantly reduced the PLA signal between Cav1-A2E and PTRF/cavin1, whereas the Cav1-A2E/filamin-A interaction was not affected (Figures 3M). Interestingly, PLA signals between Cav1-A2E and ROCK1 as well as between Cav1-A2E and HSPB1 were significantly increased upon ROCK inhibition (Figure 3N). This corroborates our proximity proteomics data and suggests that a) ROCK1 and HSPB1 are recruited to caveolae (or the cell rear) at high membrane tension, b) that ROCK activity is required for caveolae reassembly after hypo-osmotic shock, and c) that ROCK activity weakens the protein’s association with caveolae (or the cell rear).

### ARHGAP29 controls caveolin-1 phosphorylation, caveolae rear localisation and cell migration

A number of proteins identified in the Cav1 interactome had previously not been linked to caveolae (Figure 2H). One of these, the RhoGAP ARHGAP29, regulates mechanotransduction in a YAP-dependent manner and has been shown to control cancer cell migration and invasion by inhibiting actin polymerisation through a Rho/LIMK/cofilin pathway (60, 61). YAP also regulates caveolar gene transcription and caveolae formation (62). Caveolae, in turn, negatively regulate YAP (62, 63), suggesting a functional link between caveolae, YAP, and ARHGAP29-mediated Rho signaling. To test this, we downregulated ARHGAP29 or Cav1 in RPE1 cells using specific siRNAs (Figure 4A and 4B). Interestingly, ARHGAP29, YAP, and pS127 YAP protein levels were significantly increased in Cav1 siRNA cells. Moreover, we found that Y14 phosphorylation of Cav1 was elevated in ARHGAP29 siRNA cells (Figure 4A and 4B). As phosphorylation of Y14 is dependent on ROCK activity (47) this provides evidence for a function of ARHGAP29 in regulating caveolae via inhibition of Rho/ROCK signalling. Together these data suggest that caveolae dampen ARHGAP29 expression via inhibition of YAP, which promotes ROCK-dependent Cav1 Y14 phosphorylation and actomyosin-mediated contractility at the cell rear (Figure 5).

**Figure 4.**
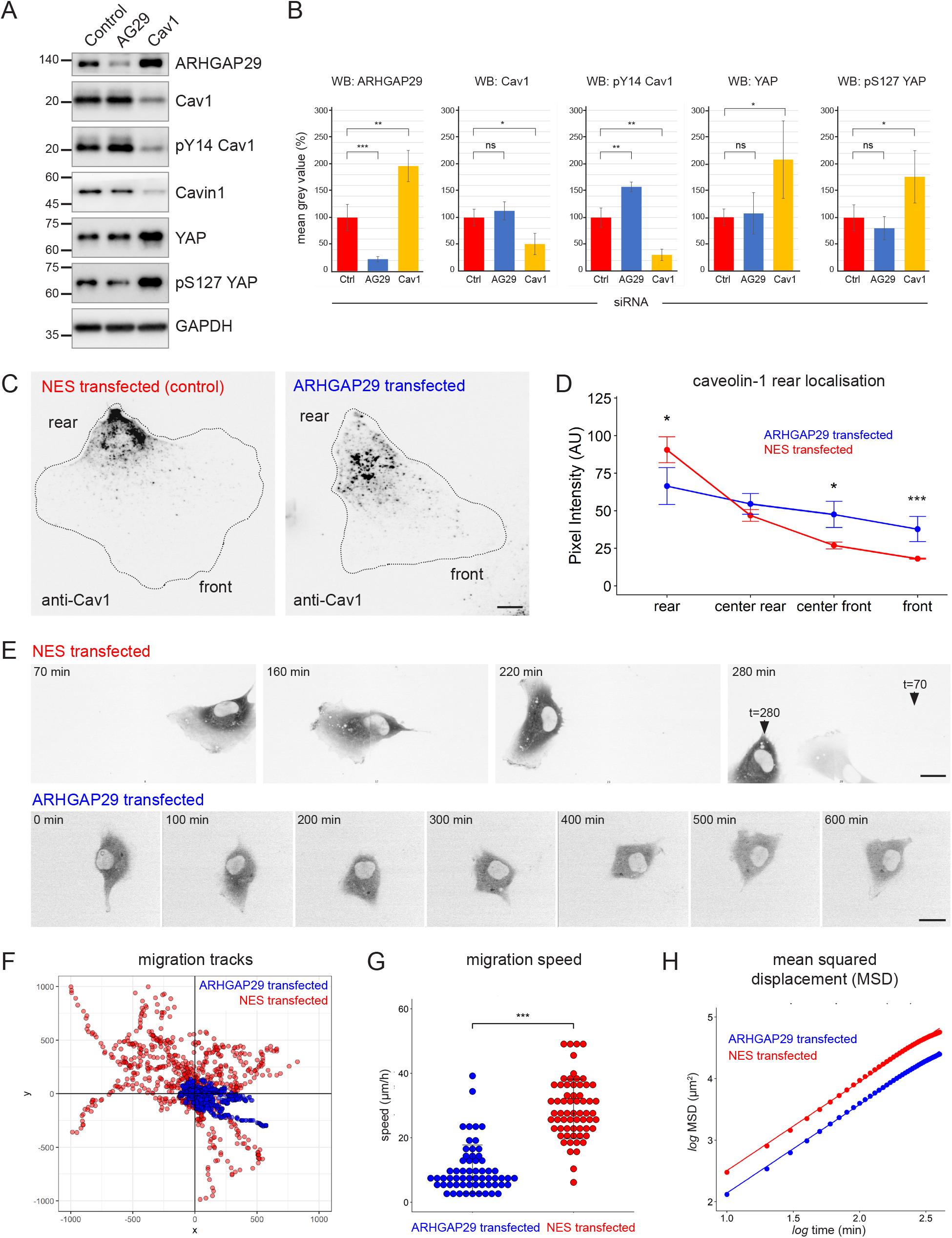
ARHGAP29 controls caveolae rear localisation and RPE1 cell migration. A) RPE1 cells were transfected with esiRNAs against ARHGAP29, caveolin-1, or non-targeting esiRNAs and analysed by Western blotting 48 hrs post-transfection. B) Densitometry analysis of Western blot band intensities as shown in A. Ratios normalized to the GAPDH loading control are displayed relative to the intensity of the control esi-RNA transfection for each protein indicated. Data represent the mean ± SD of 3-4 independent experiments. Statistical significance was calculated using an unpaired T-test. ns = p > 0.05, * = p ≤ 0.05, ** = p ≤ 0.01, *** = p ≤ 0.001. C) RPE1 cells were transfected with A2E-ARHGAP29 or NES-A2E, fixed, and stained with anti-Cav1 antibodies. D) Quantification of Cav1 rear localisation based on data shown in C. Error bars indicate mean ± SEM * = p ≤ 0.05, *** = p ≤ 0.001, Wilcoxon test (n=18 cells per condition). E) Still images of RPE1 cells transfected with NES-A2E (top) or A2E-ARHGAP29 (bottom) imaged live by spinning disk confocal microscopy. F-H) Migration tracks (F), migration speed (G) and mean squared displacement (H) of RPE1 cells transfected with A2E-ARHGAP29 or NES-A2E. Quantification was performed on three independent experiments and a total of ~60 cells per sample.

**Figure 5.**
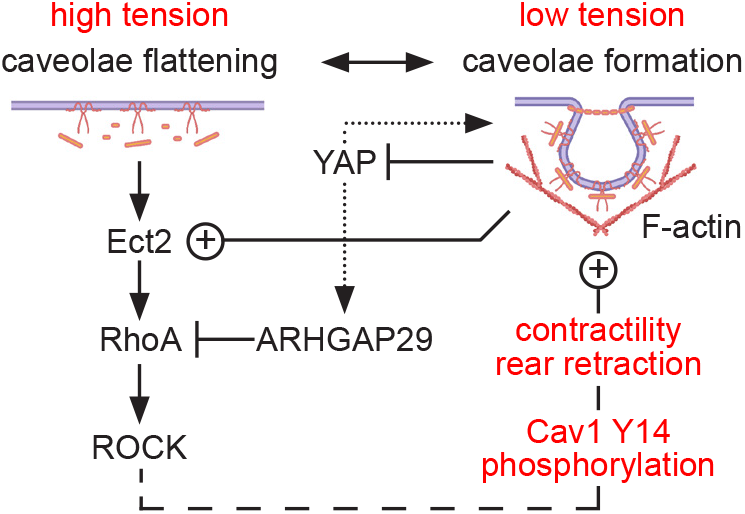
Model of ARHGAP29 function at the rear of migrating cells. ARHGAP29 blocks Rho/ROCK signalling at the cell rear, thereby counteracting the function of the RhoGEF Ect2. Caveolae block ARHGAP29 transcription via inhibition of YAP, and YAP in turn controls caveolar gene transcription and therefore caveolae formation (dotted arrows). Rho/ROCK signalling promotes Cav1 Y14 phosphorylation and cell rear contraction.

The localisation of Cav1 and PTRF/cavin1 were not notably altered in ARHGAP29 siRNA-transfected cells (data not shown). In addition, cells with reduced ARHGAP29 expression showed normal migration behaviour compared to control siRNA transfected cells (Figure S4A-D). We therefore overexpressed ARHGAP29 and analysed the subcellular distribution of Cav1 and PTRF/cavin1 in fixed cells that showed an intact actin cytoskeleton and normal cell shape. While Cav1 and PTRF/cavin1 were enriched at the rear in cells transfected with the NES-A2E control plasmid, in cells expressing moderate levels of A2E-ARHGAP29 both Cav1 and PTRF/cavin1 were displaced from the cell rear and instead localised toward the cell centre and cell front (Figure 4C and 4D and Figure S4E and S4F). We then investigated the effect of ARHGAP29 overexpression on RPE1 cell migration using time lapse microscopy (Figure 4E). Both transfected (Figure S4E) and endogenous ARHGAP29 detected with an antibody (data not shown) was localised to but not enriched at the rear of RPE1 cells. Control-transfected RPE1 cells showed stable front-rear polarity and migrated randomly but persistently, similar to non-transfected cells (Supplemental Movie S4). By contrast, cells overexpressing ARHGAP29 established multiple random protrusions and failed to specify a stable front-rear axis, resulting either in a complete loss of cell motility (Supplemental Movie S5) or in short migration tracks and reversals in direction (Supplemental Movie S6). Consequently, ARHGAP29 overexpressing cells showed markedly reduced migration velocities and mean squared displacements compared to control-transfected cells (Figure 4F–4H). Taken together these data indicated that ARHGAP29 acts as a negative regulator of caveolae rear localisation and RPE1 cell migration, most likely via inhibition of Rho-mediated actomyosin contractility.

In conclusion, using APEX2-mediated time-resolved proximity proteomics we have identified a dynamic Cav1 interactome that is highly sensitive to acute changes in membrane tension. We propose that this protein network regulates caveolae rear localisation, rear retraction and mechanotransduction in migrating cells. Previous studies indicate a role for Rho/ROCK signalling and the RhoGEF Ect2 in caveolae formation and cell rear retraction (39, 40). We show here that ROCK is recruited to Cav1 at high membrane tension (i.e. under conditions that promote caveolae flattening) and that ROCK activity is required for caveolae reassembly after hypo-osmotic shock. Conversely, the RhoGAP ARHGAP29 is associated with caveolae at low membrane tension and inhibits caveolae rear localisation and cell migration when overexpressed. This suggests that Ect2/ROCK and ARHGAP29 have apposing functions in caveolae formation and in fine-tuning actomyosin contractility at the cell rear (Figure 5). How exactly caveolae regulate ROCK, Ect2 and ARHGAP29, and how their activities in turn control cycles of caveolae formation and disassembly remains to be explored. It is equally unclear at present how these proteins are recruited to the cell rear. ARHGAP29 contains an N-terminal F-BAR domain that can sense changes in membrane curvature (64). Subtle fluctuations in membrane tension and curvature may therefore permit ARHGAP29 to cycle on and off the rear membrane. Membrane binding of ARHGAP29 is expected to temporally dampen RhoA-mediated contractility, causing a transient increase in rear membrane tension and caveolae flattening/disassembly. Cycles of caveolae assembly and disassembly could thereby coordinate rear retraction with protrusive forces at the cell front, permitting persistent cell migration. Taken together our data and previous work (39, 40) suggest that caveolae act as a rear-localised membrane tension sensor that is critical for linking actomyosin contractility to cell rear retraction. The caveolar interactome identified here will be instrumental in dissecting the molecular machinery and mechanisms underlying caveolae-mediated mechanotransduction in migrating cells.

## Supporting information

TableS1_Protein Interactions

Key Resource Table

## Acknowledgements

RG and EM were supported by the FNR AFR Bilateral Singapore grant 11823257 to GD and AL. Illustrations were made using Biorender.

## Author contributions

RG made cell lines, generated and analysed the proteomics data, performed and analysed the PLA assays, and prepared the figures. EM performed all time-lapse imaging experiments, cell migration assays, and the siRNA experiments. AL performed all EM experiments and wrote the manuscript. AL and GD supervised RG and EM and conceived the study.

## Data availability

The raw mass spectrometry data is available through the PRIDE repository, ID: PXD026464.

## Materials and Methods

### DNA constructs and bacterial strains

Cav1-APEX2-EGFP and NES-APEX2-EGFP were previously described (7, 65). To produce EHD2-A2E, human EHD2 cDNA was cloned into the N1-A2E vector (65) using the *HindIII* and *XhoI* sites. To produce EGFP-APEX2-ARHGAP29 the ARHGAP29 cDNA was sub-cloned into the C1-A2E vector (65) by PCR using HA-ARHGAP29 (Addgene plasmid #104154) as a template. All primers and oligonucleotides are listed in the Materials table. NEB10-beta competent *E. coli* (NEB, Cat#C3019I) were transformed for plasmid amplification and then cultured in LB Agar. Single colonies were isolated and further expanded in LB broth before plasmid purification with miniprep (Qiagen) following the manufacturer protocol. All plasmids were verified by Sanger sequencing.

### RPE1 cell culture and transfection

RPE1 cells were cultured in DMEM-F12 supplemented with 10% FBS, 100 units/mL penicillin, and 100 μg/mL streptomycin (GIBCO) at 37°C and 5% CO_2_. Transfections were performed in 6-well plates two days post-seeding with the cells in a sub-confluent condition. Cells were transfected with Lipofectamine-3000 (Thermo Fisher Scientific) using 1 μg plasmid DNA per well. Stable cell lines were generated by selection in complete medium containing 400 μg/ml geneticin (Gibco) for at least 14 days. Stable cell lines were maintained in media containing 200 μg/ml geneticin. Cells were tested regularly for mycoplasma using a PCR kit.

### Hypo-osmotic shock and inhibitors

For hypo-osmotic shock cells were shifted from normal growth medium to hypo-osmotic medium (10% DMEM/90% deionized water) for 5 min and then immediately collected for the Hypo-osmotic condition, or switched back to normal medium for 30 min to let cells recover from the osmotic shock. For the ROCK inhibition assays, the 5 min shock was performed in the presence or absence of 10 μM Y27632 (DMSO (1 μl/ml) was used as a control), followed by 30 min recovery in the presence or absence of the inhibitor.

### Immunofluorescence

RPE1 cells grown on fibronectin-coated glass coverslips were washed twice in PBS and fixed in 4% paraformaldehyde (PFA) for 15 min. After further washing, cells were permeabilised with 0.1% Triton X-100 in PBS for 5 min. Blocking was performed with 10% FBS in PBS for 1h at RT. Samples were incubated for 1h at RT with primary antibodies diluted in 0.1% BSA, 0.01% Tween-20 in PBS (antibody buffer). After 3 washes with antibody buffer, samples were further incubated with the appropriate fluorescent-conjugated secondary antibody for 1h RT. After 3 last washes in PBS, coverslips were mounted in Vectashield Antifade Mounting Medium containing DAPI (Vector Laboratories). Slides were imaged either on a spinning disk microscope (CorrSight, Thermo Fisher Scientific) equipped with an Orca R2 CCD camera (Hamamatsu) or on a laser scanning confocal microscope (LSM880 FastAiry, Carl Zeiss). Images were acquired with a 63× oil immersion Plan-Apochromat objective (NA=1.4, Zeiss) and standard filter sets. Confocal stacks were processed in ImageJ/FIJI.

### Electron Microscopy

For electron microscopy cells were fixed in 2.5%glutaraldehyde (EM grade, EMS) in 0.1 M cacodylate buffer pH 7.4 (CB) for 1 h. After several washes in CB cells were incubated in 0.5 mM diaminobenzidine free-base (DAB, Sigma-Aldrich) and 0.5 mM H_2_O_2_ in CB for 5-10 min, post-fixed in 1% osmium tetroxide (EMS), 1% (v/w) potassium ferricyanide (Sigma-Aldrich) in CB, dehydrated using a graded ethanol series, and further processed for TEM as described previously (66). For CLEM areas containing the cells of interest were sawed out using a jeweller’s saw and sectioned parallel to the substratum using a diamond knife. 70-80 nm semithin serial sections were picked up on formvar- and carbon-coated EM slot grids and imaged on a TecnaiT12 TEM (Thermo Fisher Scientific) operated at 120 kV equipped with a 4k × 4k Eagle camera (Thermo Fisher Scientific). MiniSOG labeling and 3D tomography was described previously (8). Tomograms were reconstructed using weighted back projection and further analysed in IMOD/etomo.

### Proximity ligation assay

*In situ* PLA was performed with DuoLink PLA kit (Sigma-Aldrich) following the manufacturer’s protocol. Briefly, after fixation and permeabilization steps described above, cells were first incubated in Duolink Blocking Solution for 1h at 37°C and then with the appropriate mix of primary antibodies for 1h RT. After washes, coverslips were incubated with PLA probe solution for 1h at 37°C. Ligation and amplification steps were performed at 37°C for 30 and 100 min, respectively. Coverslips were mounted with Duolink PLA Mounting Medium with DAPI.

### PLA Imaging and Quantification

Slides were imaged on a Zeiss LSM 880 confocal microscope and images were acquired with a 63× oil immersion Plan-Apochromat objective 1.4 numerical aperture (Zeiss) and standard filter sets. Image analysis was performed with ImageJ/FIJI according to published protocols. Briefly, single stack images were split in separate channels. The green channel was used for cell boundaries identification while the red channel for PLA signal retrieval. 30 different cells from three independent experiments were used for quantification. The dot plots represent the number of PLA signals per cell.

### Proximity biotinylation

The APEX2 labelling protocol was adapted from previously published papers (44, 46, 65). Briefly, cells were grown on glass coverslips (for IF) or on plastic dishes (for MS and WB). Cells were incubated for 30 min at 37°C in media containing 0.5 mM biotin phenol (Iris Biotech). After 2 washes in PBS, the labelling reaction was performed for 1 min with 0.5 mM H_2_O_2_ in PBS. Due to the higher expression level of NES-A2E compared to Cav1-A2E, we adjusted the hydrogen peroxide concentration to obtain an overall biotinylation intensity comparable to that obtained in the Cav1-A2E cell line. To quench the reaction, three washes with a quencher solution (5 mM (±)-6-hydroxy-2,5,7,8-tetramethylchromane-2-carboxylic acid (Trolox), 10 mM sodium ascorbate, 10 mM sodium azide in PBS) were performed. Cells were then further processed either for IF or protein extraction.

### Protein extraction and Western blotting

Cells were incubated in lysis buffer (1% SDC in 50mM Ammonium Bicarbonate) and scraped using a cell scraper. Lysates were then sonicated (three cycles of pulse sonication for 20 seconds) and clarified (centrifugation at 16,000xg for 30 min at 4°C). Protein concentration was assessed either with Bradford’s assay or with EZQ protein quantification assay (Thermo Fisher Scientific) and then kept at −80°C until further processing. Proteins were separated on precast 4-12% polyacrylamide gels and blotted on polyvinylidene difluoride (PVDF) membranes. Membranes were then dehydrated in methanol for 10 s and allowed to dry for 1 h. For anti-Cav1 and anti-GFP, membranes were incubated with antibodies (anti-Cav1 1:10,000; anti-GFP 1:1,000) diluted in 2% bovine serum albumin (BSA) in PBST (PBS, 0.2% Tween-20). After 3×10 min washes in PBST, membranes were incubated with HRP-conjugated secondary antibody diluted in 5% fat-free milk in PBST. After further washes, membranes were developed with chemiluminescent substrate. To test biotinylation efficiency, membranes were incubated with streptavidin-HRP conjugated diluted 1:10,000 in PBST containing 2% BSA for 1h at RT and then washed 5 times with PBST before development.

### Streptavidin purification and on-bead digestion

Streptavidin Beads (Pierce™ Streptavidin magnetic beads, Thermo Fisher Scientific) were washed twice with cold lysis buffer before incubation with protein lysates (50 μl bead slurry per 1 mg lysate) for 1.5h at 4°C on a rotating wheel. Beads were then washed twice with cold lysis buffer, and then once with each of the following washing buffers: 1M KCl, 0.1M of Na_2_CO_3_, 2M Urea in 50 mM Ammonium bicarbonate (AmBic) pH 8. The last two washes were performed with cold 50 mM AmBic pH 8. Beads were then re-suspended in digestion buffer (2M Urea in 50 mM AmBic). Protein reduction was performed by adding 1 M DTT to reach 10 mM final concentration, and incubating for 45 min at 37°C and 800 rpm. After incubation with Iodocaetamide (25 mM final concentration, 30 min at RT and 800 rpm), alkylation reaction was stopped by incubating the samples at natural light for 2 min. Protein digestion was performed in a 2-step reaction: samples were pre-digested with 0.5 μg LysC for 4h at 37°C and 800 rpm and then digested with 1 μg trypsin O/N at 37°C and 800 rpm. Peptides were then acidified (pH 2) by adding Formic Acid to 1% final concentration and centrifuged for 15 min at 16000 rpm at 4°C to remove any residues of beads before proceeding with desalting on Sep-Pak μ-C18 Elution Plates. Peptides were then dried on a speed vac for 2 hours and re-suspended in 0.1% formic acid prior to LC-MS.

### Mass spectrometry

LC-MS/MS was performed using an Ultimate 3000 RSLCnano (Thermo Fisher Scientific) LC system equipped with an Acclaim PepMap RSLC column (15 cm × 75 μm, C18, 2 μm, 100 Å, Thermo Fisher Scientific). Peptides were eluted at a flow rate of 300nL/min in a linear gradient of 2-35% solvent B over 70 min (solvent A: 0.1% formic acid; B: 0.1% formic acid in acetonitrile (ACN)), followed by a 5 min washing step (90% solvent B) and a 10 min equilibration step (2% solvent B). Mass spectrometry was performed on a Q-Exactive Plus mass spectrometer (Thermo Fisher Scientific) equipped with a nano-electrospray source and using uncoated SilicaTips (12cm, 360μm o.d., 20μm i.d., 10μm tip i.d.) for ionization, applying a 1500 V liquid junction voltage at 250°C capillary temperature. MS/MS analysis were performed in data dependent acquisition (DDA) mode, and the 12 most intense parent ions were fragmented with a resolving power of 17500 (at 200m/z). Automatic gain control (AGC), maximum fill time and dynamic exclusion were set to 1e6, 60 ms and 30 ms, respectively.

### Data analysis for mass spectrometry

Protein quantification was performed with MaxQuant (version 1.6.7.0) using the following parameters: carbamidomethyl cysteine as fixed modification, methionine oxidation and N-acetylation as variable modifications, digestion mode trypsin specific with maximum 2 missed cleavages, and an initial mass tolerance of 4.5 ppm for precursor ions and 0.5 Da for fragment ions. All experiments were performed in triplicates and the abundance of assembled proteins was determined using label-free quantification (standard MaxQuant parameters) and match between run option was selected. Protein identification was performed searching against the Uniprot reference proteome (Uniprot ID 9606, downloaded on 06 January 2020). To this database were added 2 sequences for the detection of the fusion protein: Cav1-CANLF (Uniprot id: P33724) and a manually curated fusion protein combining APEX2 (Q43758) and EGFP (P42212). Further processing with Perseus (version 1.6.10.45) was performed according to the following pipeline: removal of common contaminants, reverse and peptides identified only on site-specific modifications, log2 transformation. Only proteins identified in 3 out of 3 replicates of at least 1 sample were considered for further analysis. Moderated t-test was performed with Proteomics Toolset for Integrative Data Analysis (ProTIGY, Broad Institute, Proteomics Platform, https://github.com/broadinstitute/protigy) and a p-value<0.05 was considered statistically significant. Further data analysis and visualization were performed in R (version 4.0.3).

### Generation of the Cav1 interactome

Interactions among the proteins identified in the Cav1 proximity proteome were retrieved from BioGRID version 4.2.193 (April 2020) and the STRING database. For BioGRID entries only physical protein interactions were considered; methods such as Affinity Capture-RNA, Protein-RNA, and Proximity Label-MS were disregarded. For STRING the following parameters were used: Active interaction; source: Database; Confidence: > 0.4. In addition, direct or indirect protein-protein interactions reported in the primary literature were included (see Table S1). The protein network was drawn in Cytoscape version 3.7.1.

### Live cell imaging and cell migration analysis

RPE1 cells were grown on fibronectin-coated glass-bottom dishes (MatTec Corp., Ashland, MA) and imaged live in complete DMEM without Phenol Red on a CorrSight spinning disk microscope (Thermo Fisher Scientific). Imaging was carried out at 37°C, 5% CO_2_ and 90% humidity in a closed atmosphere chamber (IBIDI). Up to 5 different locations were imaged simultaneously using the multi-stage position function in LA software (Thermo Fisher Scientific). Confocal images were acquired as 3×3 tiles with a 40x oil objective (NA 1.3, EC Plan Neofluar M27, Zeiss) using the 488 nm laser line (65 mW; iChrome MLE-LFA) and a standard GFP filter set. Focus was maintained by a hardware autofocus system (Focus Clamp). The laser output power and exposure times were set to a minimum. Time lapse recordings were analyzed in ImageJ/FIJI (67, 68). Cell migration data were quantified in R with the CelltrackR package.

### Quantification of rear localisation index

Transfected RPE1 cells were seeded on glass coverslips at low density, fixed, and stained for Cav1, cavin1, and phalloidin. Quantification of Cav1 distribution was performed on 16-bit images in ImageJ/FIJI by drawing a line (50 pixel wide) from the rear to the front, along the longest cell axis. The pixel intensity distribution along the line was measured using the Plot Profile function. Background was subtracted and the pixel intensities along the line were divided into four equal size quadrants - rear, center rear, center front, and front. The average pixel intensities in each quadrant were determined and plotted.

### siRNA transfection

esiRNAs specific to human caveolin-1 or ARHGAP29 and a non-targeting esiRNAs were purchased from Sigma-Aldrich and transfected at a final concentration of 50 nM using Dharmafect (Sigma-Aldrich) according to the manufacturer’s recommendations. Transfected cells were analysed by Western blotting 48 hrs post-transfection. For immunofluorescence and cell migration analysis, cells were trypsinized 48 hrs post-transfection and seeded at low density onto glass coverslips or MatTec dishes, respectively. Cells were imaged 24 hrs later.

