## Supplementary Material for "Time-resolved proximity proteomics uncovers a membrane tension-sensitive caveolin-1 interactome at the rear of migrating cells"

**Figure S1: Characterisation of RPE1 cell lines**

A) Western blot analysis of protein lysates from RPE1 cells stably expressing Cav1-APEX2-EGFP or NES-APEX2-EGFP fusion proteins. The membrane was probed with anti-GFP antibodies.

B) Representative immunofluorescence stainings of Cav1-A2E RPE1 cells at isotonic (NT) and hypotonic (HYPO) conditions, and after recovery from hypo-osmotic shock (REC). Cells were co-stained with anti-PTRF/cavin1 antibodies.

C) $H_2O_2$ ramp of NES-APEX2-EGFP expressing RPE1 cells pre-incubated for 30 min with 0.5 mM biotin phenol. Increasing concentrations of $H_2O_2$ were used during the biotinylation reaction to determine the optimal $H_2O_2$ concentration for further experiments.

B) Confocal fluorescence microscopy of NES-APEX2-EGFP cells after proximity labelling with APEX2. Cells were fixed and stained with fluorescent streptavidin (Alexa Fluor 555) to visualize biotinylated proteins. Nuclei were counterstained with DAPI. Scale bars 20 µm.

**Figure S2: Quantitative proximity proteomics of Cav1-APEX2-EGFP in response to hypo-osmotic shock.** Heatmap of 111 z-scored LFQ quantified protein groups differentially enriched across the three treatments (non-treated (NT, control), hypo-osmotic shock (HYPO), and recovery from hypo-osmotic shock (REC). All three replicates are shown. NA indicates not identified.

**Figure S3: Proximity ligation assays in RPE1 cells**

A, C, E) Representative images of Proximity Ligation Assays on Cav1-A2E expressing RPE1 cells using anti-GFP antibodies and either anti-FLNA (A), anti-CTTN (C), or anti-HSPB1(E) antibodies. GFP and DAPI signals used to determine cell boundaries are shown in the upper panels, while the bottom panels show the PLA signal masks used for counting. Scale bars: 20 µm.

B,D,F,H) Representative immunofluorescence stainings of Cav1-A2E cells. The upper panel shows the GFP signal, while the bottom panels show anti-FLNA (B), anti-CTTN (D), anti-HSPB1 (F) and anti-ROCK1 (H) signals.

G) PLA negative control. Samples were probed only with anti-GFP primary antibody and both PLA probes.

**Figure S4: Overexpression and siRNA-mediated knockdown of ARHGAP29**

A-D) Migration tracks (A), migration speed (B), displacement (C), and mean squared displacement (D) of RPE1 cells transfected with esiRNAs against ARHGAP29 or non-targeting esiRNAs. Quantification was performed on two independent experiments.

E) RPE1 cells transfected with A2E-ARHGAP29 were fixed and stained with anti-Cavin1 antibodies and fluorescently labelled phalloidin (Alexa-Fluor 555). Asterisks (*) indicate two untransfected control cells. White arrowheads indicate enrichment of cavin1 at the cell rear. Red arrowheads indicate actin and ARHGAP29 at the cell rear.

F) 2D and 3D heat maps of two untransfected control cells (asterisks in A) and an A2E-ARHGAP29 transfected cells. White arrowheads indicate the position of the cell rear. Note the loss of cavin1 enrichment at the cell rear in cells transfected with A2E-ARHGAP29.

**Supplemental Movie S1:** Time lapse microscopy of an RPE1 cell stably expressing cavin3-miniSOG-mCherry migrating on fibronectin-coated glass imaged by spinning disk microscopy.

**Supplemental Movie S2:** Time lapse microscopy of an RPE1 cell stably expressing cavin3-miniSOG-mCherry migrating in a 3D collagen matrix imaged by spinning disk microscopy.

**Supplemental Movie S3:** Electron tomogram of caveolar clusters and networks at the rear of RPE1 cells stably expressing cavin3-miniSOG-mCherry. Caveolae were specifically labelled using miniSOG.

**Supplemental Movie S4:** Time lapse microscopy of RPE1 cells transfected with an NES-APEX2-EGFP plasmid imaged by spinning disk microscopy.

**Supplemental Movie S5:** Time lapse microscopy of RPE1 cells transfected with an APEX2-EGFP-ARHGAP29 plasmid imaged by spinning disk microscopy.

**Supplemental Movie S6:** Time lapse microscopy of RPE1 cells transfected with an APEX2-EGFP-ARHGAP29 plasmid imaged by spinning disk microscopy.

### Figure S1

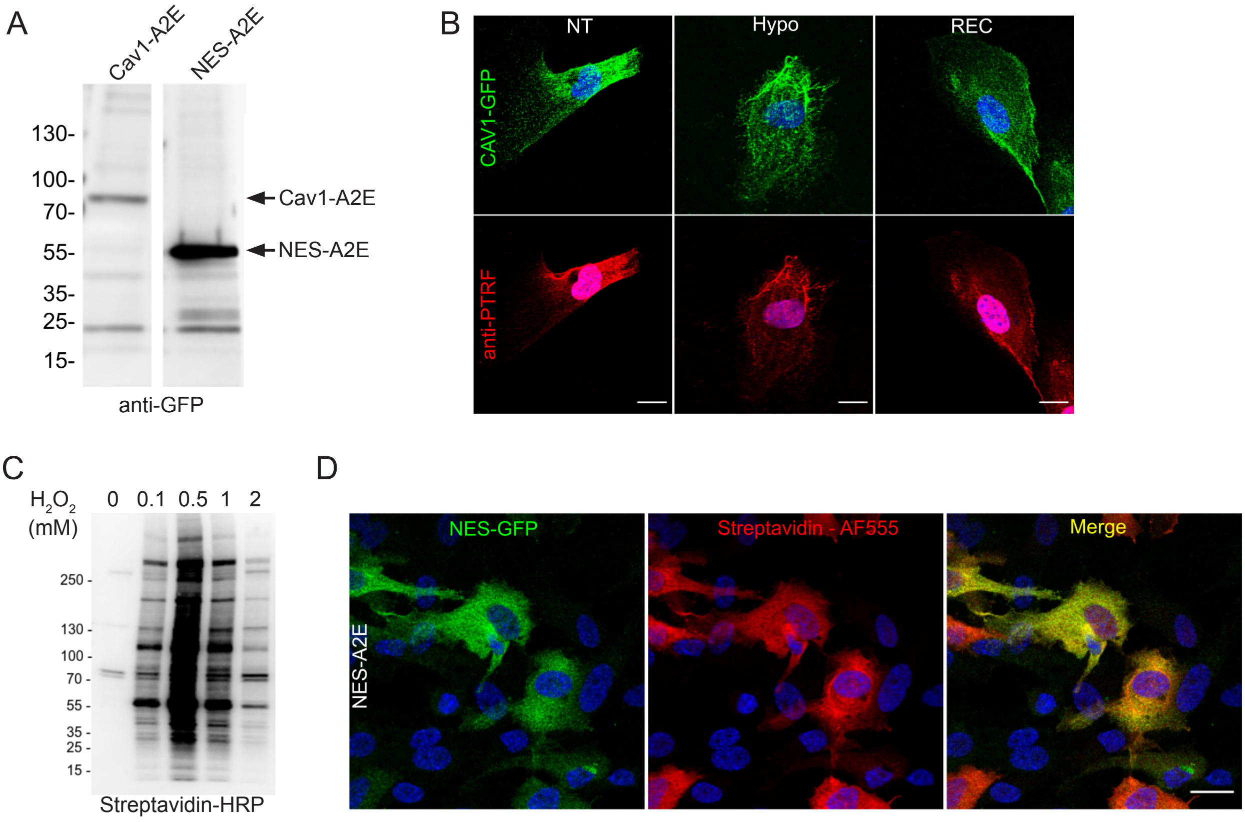

A
Cav1-A2E
NES-A2E
130-
100-
70-
55-
35-
25-
15-
Cav1-A2E
NES-A2E
anti-GFP
B
NT
Hypo
REC
CAV1-GFP
anti-PTRF
C
H2O2 (mM)
0
0.1
0.5
1
2
250 -
130 -
100 -
70 -
55 -
35 -
25 -
15 -
Streptavidin-HRP
D
NES-GFP
Streptavidin - AF555
Merge
NES-A2E

### Figure S2

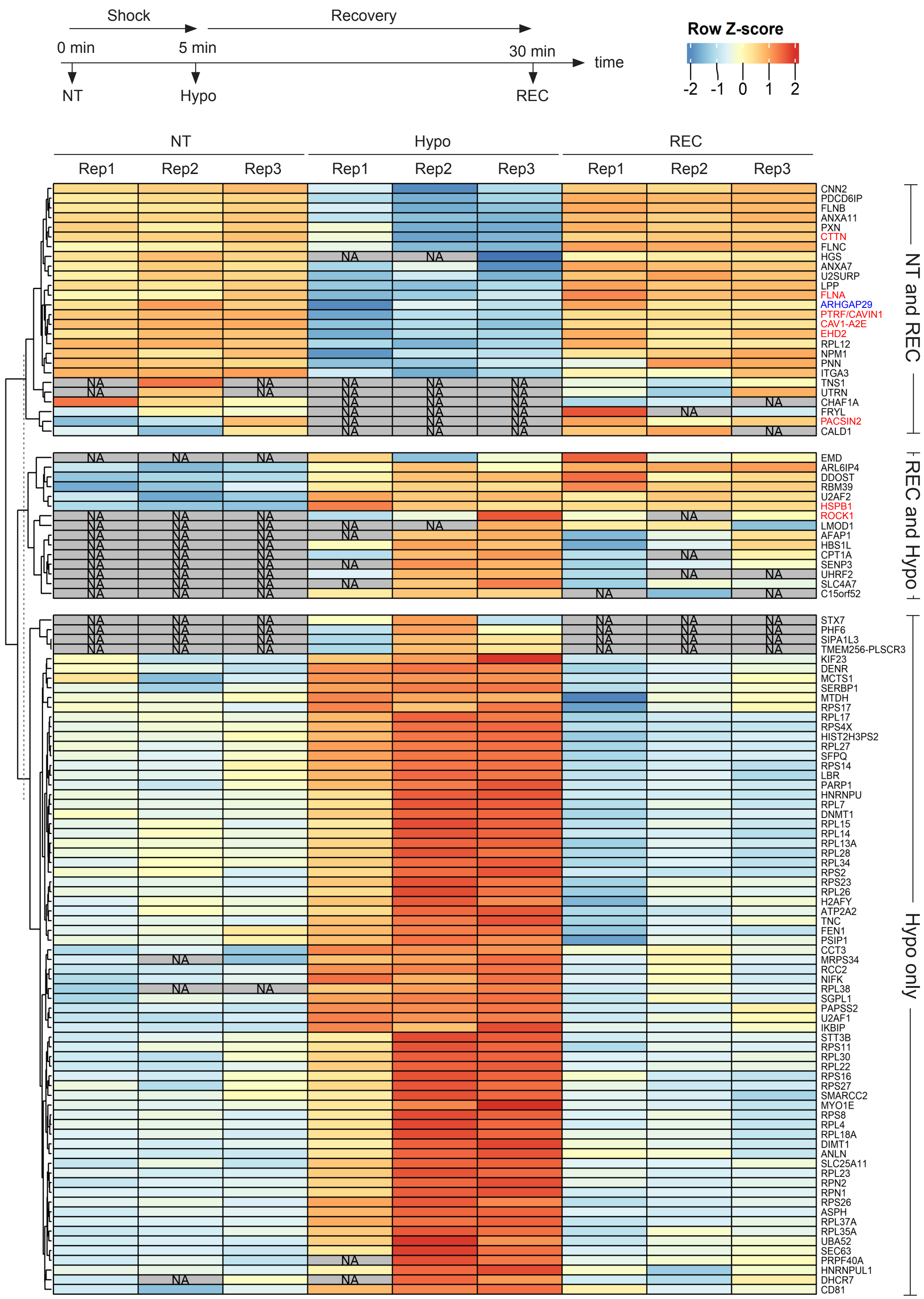
Shock
Recovery
0 min
5 min
30 min
time
NT
Hypo
REC
Row Z-score
-2 -1 0 1 2
NT
Hypo
REC
Rep1 Rep2 Rep3 Rep1 Rep2 Rep3 Rep1 Rep2 Rep3
NA
CNN2
PDCD6IP
FLNB
ANXA11
PXN
CTTN
FLNC
HGS
ANXA7
U2SURP
LPP
FLNA
ARHGAP29
PTRF/CAVIN1
CAV1-A2E
EHD2
RPL12
NPM1
PNN
ITGA3
TNS1
UTRN
CHAF1A
FRYL
PACSIN2
CALD1
NT and REC
EMD
ARL6IP4
DDOST
RBM39
U2AF2
HSPB1
ROCK1
LMOD1
AFAP1
HBS1L
CPT1A
SENP3
UHRF2
SLC4A7
C15orf52
REC and Hypo
STX7
PHF6
SIPA1L3
TMEM256-PLSCR3
KIF23
DENR
MCTS1
SERBP1
MTDH
RPS17
RPL17
RPS4X
HIST2H3PS2
RPL27
SFPQ
RPS14
LBR
PARP1
HNRNPU
RPL7
DNMT1
RPL15
RPL14
RPL13A
RPL28
RPL34
RPS2
RPS23
RPL26
H2AFY
ATP2A2
TNC
FEN1
PSIP1
CCT3
MRPS34
RCC2
NIFK
RPL38
SGPL1
PAPSS2
U2AF1
IKBIP
STT3B
RPS11
RPL30
RPL22
RPS16
RPS27
SMARCC2
MYO1E
RPS8
RPL4
RPL18A
DIMT1
ANLN
SLC25A11
RPL23
RPN2
RPN1
RPS26
ASPH
RPL37A
RPL35A
UBA52
SEC63
PRPF40A
HNRNPUL1
DHCR7
CD81
Hypo only

### Figure S3

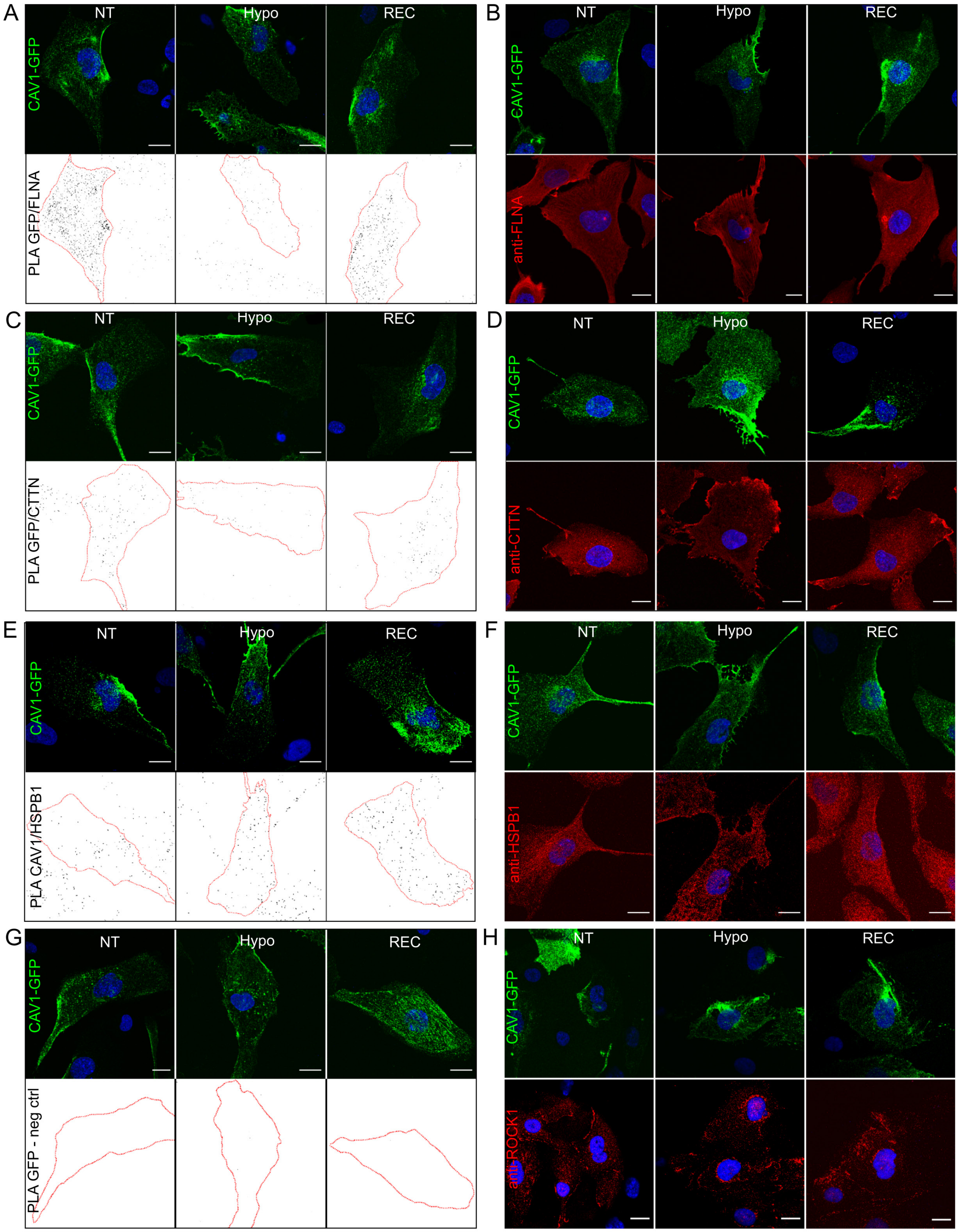
A
NT
Hypo
REC
CAV1-GFP
PLA GFP/FLNA
B
NT
Hypo
REC
CAV1-GFP
anti-FLNA
C
NT
Hypo
REC
CAV1-GFP
PLA GFP/CTTN
D
NT
Hypo
REC
CAV1-GFP
anti-CTTN
E
NT
Hypo
REC
CAV1-GFP
PLA CAV1/HSPB1
F
NT
Hypo
REC
CAV1-GFP
anti-HSPB1
G
NT
Hypo
REC
CAV1-GFP
PLA GFP - neg ctrl
H
NT
Hypo
REC
CAV1-GFP
anti-ROCK1

### Figure S4

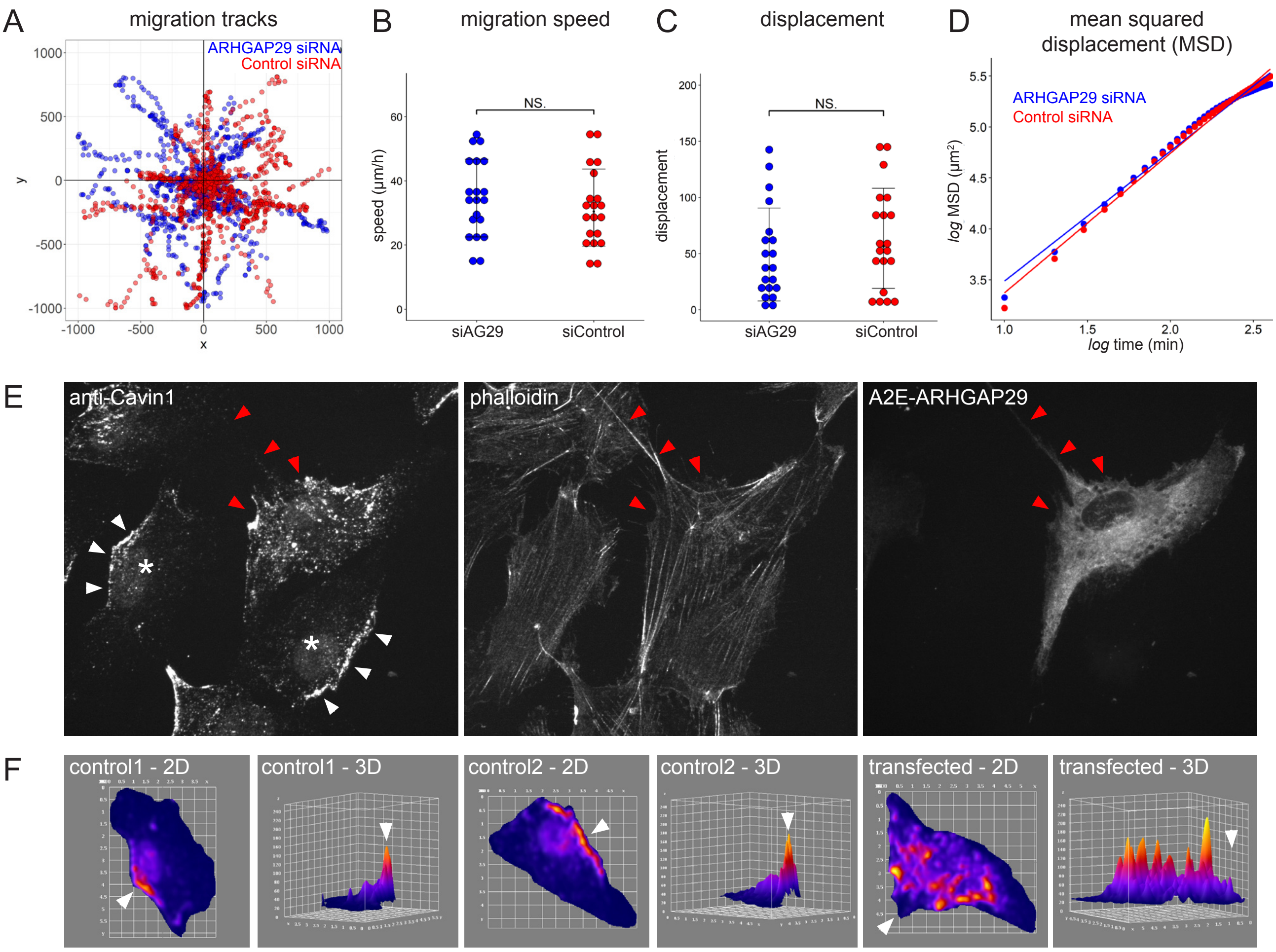
A
migration tracks
ARHGAP29 siRNA
Control siRNA
B
migration speed
NS.
speed (µm/h)
siAG29
siControl
C
displacement
NS.
displacement
siAG29
siControl
D
mean squared displacement (MSD)
ARHGAP29 siRNA
Control siRNA
log MSD (µm²)
log time (min)
E
anti-Cavin1
phalloidin
A2E-ARHGAP29
F
control1 - 2D
control1 - 3D
control2 - 2D
control2 - 3D
transfected - 2D
transfected - 3D
