## Supplementary material for "Time-resolved proximity proteomics uncovers a membrane tension-sensitive caveolin-1 interactome at the rear of migrating cells": Key Resource Table

| REAGENT or RESOURCE | SOURCE | IDENTIFIER |
| --- | --- | --- |
| *Cell lines and Cell culture* | | |
| hTERT RPE-1 | ATCC | ATCC CRL-4000 |
| DMEM:F12 1:1 Mixture with Ultraglutamine I, 500ml | Westburg | Cat#LO BE04-687F/U1 |
| PBS (1X) without Ca++, Mg++, 500ml | Westburg | Cat#LO BE17-516F |
| Trypsin/Versene(EDTA) (1X ) 100mL | Westburg | Cat#LO BE17-161E |
| Gibco™ Penicillin-Streptomycin (10,000 U/mL) | Fisher Scientific | Cat#15140122 |
| Geneticin™ Selective Antibiotic (G418 Sulfate) (50 mg/mL) | Fisher Scientific | Cat#10131027 |
| Gibco™ Fibronectin Bovine Protein, Plasma | Fisher Scientific | Cat#33010018 |
| Lipofectamine 3000 Reagent | Thermo Fisher Scientific | Cat# L3000015 |
| *Antibodies and PLA reagents* | | |
| Rabbit monoclonal anti-PARG1 (ARHGAP29) | Invitrogen | Cat#PA5-55336 |
| Rabbit polyclonal anti-Caveolin-1 (CAV1) | BD Bioscience/Transduction labs | Cat#610060 |
| Mouse monoclonal anti-Caveolin-1 pY14 | BD Bioscience/Transduction labs | Cat#611339 |
| Rabbit polyclonal anti-Cavin-1 (PTRF) | Abcam | Cat#ab48824 |
| Goat polyclonal anti-EHD2 | Abcam | Cat#ab23935 |
| Mouse monoclonal anti-Filamin 1 (E-3) (FLNA) | Santa Cruz Biotechnology | Cat#sc-17749 |
| Mouse monoclonal anti-Cortactin (clone 4F11) | Merck Millipore | Cat#05-180 |
| Mouse monoclonal anti-Rock-1 (G-6) | Santa Cruz Biotechnology | Cat#sc-17794 |
| Mouse monoclonal anti-HSP27 (F-4) (HSPB1) | Santa Cruz Biotechnology | Cat#sc-13132 |
| Mouse monoclonal anti-GFP | Roche | Cat# 11814460001 |
| Rabbit polyclonal anti-GFP | Abcam | Cat#ab290 |
| Mouse monoclonal anti-YAP | Cell Signalling Technology | Cat #14074 |
| Mouse monoclonal anti-YAP pS127 | Cell Signalling Technology | Cat #13008 |
| Donkey anti-Mouse IgG (H+L) Alexa Fluor 555 | Invitrogen | Cat# A-31570 |
| Donkey anti-Rabbit IgG (H+L) Alexa Fluor 555 | Invitrogen | Cat# A-31572 |
| Goat anti-Rabbit IgG (H+L) Alexa Fluor 633 | Invitrogen | Cat# A-21071 |
| Goat anti-Mouse IgG (H+L) Alexa Fluor 633 | Invitrogen | Cat# A-21052 |
| Goat anti-rabbit IgG (H+L) HRP conjugate | Invitrogen | Cat# A16104 |
| Goat anti-mouse IgG (H+L) HRP conjugate | Invitrogen | Cat# A16072 |
| Streptavidin Alexa Fluor 568 | Thermo Fisher Scientific | Cat#S11226 |
| Duolink™ In Situ Orange Starter Kit Mouse/Rabbit | Sigma-Aldrich | Cat#DUO92102-1KT |
| *Chemicals, Peptides, and Recombinant Proteins* | | |
| Biotin phenol | Iris Biotech | Cat# LS-3500 |
| Trolox | Sigma-Aldrich | Cat# 238813 |
| Glutaraldehyde (32%) | Electron Microscopy Sciences (EMS) | Cat# 16220 |
| Paraformaldehyde (32%) | Electron Microscopy Sciences (EMS) | Cat# 100504-858 |
| Diaminobenzidine (DAB) (Free-Base) | Sigma-Aldrich | Cat# D8001 |
| Durcupan ACM resin | Electron Microscopy Sciences (EMS) | Cat# 44610 |
| Osmium Tetroxide (2%) | Electron Microscopy Sciences (EMS) | Cat# 19152 |
| Potassium Ferricyanide | Sigma-Aldrich | Cat# 702587 |
| Sequencing Grade Modified Trypsin | Promega Corporation | Cat#V5111 |
| Lysyl Endopeptidase, MS Grade | FUJIFILM Wako Pure Chemical Corporation | Cat#125-05061 |
| cOmplete™, EDTA-free Protease Inhibitor Cocktail | Roche | Cat# 11873580001 |
| Thermo Scientific™ Pierce™ Streptavidin Magnetic Beads | Fisher Scientific | Cat#10615204 |
| EZQ™ Protein Quantitation Kit | Fisher Scientific | Cat#R33201 |
| VECTASHIELD Antifade Mounting Medium with DAPI | Vector Laboratories | Cat#H-1200-10 |
| Plasmids, Oligonucleotides, and esiRNAs | | |
| Caveolin-1 siRNA ON-TARGET Plus SMART pool | Dharmacon | Cat#L-003467-00-0020 |
| ARHGAP29 Mission esiRNA:  GCTTCCCTTGCAGACAGTTTACAGTCTCTCTGTGATAGTGCCAAACTCTATGACCCAGGCCAAGAGTACAGTGAATTTGTCAAGGCCACAAATTCAACTGAAGAAGAAAAAGTTGATGGAAATGTAAATAAACATTTAAATAGTTCCCAACCTTCAGGATTTGGACCTGCCAACTCTTTAGAGGATGTTGTACGCCTTCCTGACAGTTCTAATAAAATTGAAGAGGACAGATGCTCTAACAGTGCAGATATAACAGGTCCTTCCTTTATAAGATCATGGACATTTGGGATGTTTAGTGATTCTGAGAGCACTGGAGGGAGCAGCGAATCTAGATCTCTGGATTCAGAATCTATAAGTCCAGGAGACTTTCATCGAAAACTTCCACGAACACCATCCAGTGGA | Sigma | Cat#EHU13457 |
| FLUC Control Mission esiRNA:  GAGCAACTGCATAAGGCTATGAAGAGATACGCCCTGGTTCCTGGAACAATTGCTTTTACAGATGCACATATCGAGGTGGACATCACTTACGCTGAGTACTTCGAAATGTCCGTTCGGTTGGCAGAAGCTATGAAACGATATGGGCTGAATACAAATCACAGAATCGTCGTATGCAGTGAAAACTCTCTTCAATTCTTTATGCCGGTGTTGGGCGCGTTATTTATCGGAGTTGCAGTTGCGCCCGCGAACGACATTTATAATGAACGTGAATTGCTCAACAGTATGGGCATTTCGCAGCCTACCGTGGTGTTCGTTTCCAAAAAGGGGTTGCAAAAAATTTTGAACGTGCAAAAAAAGCTCCCAATCATCCAAAAAATTATTATCATGGATTCTAAAACGGATTACCAGGGATTTCAGTCGATGTACACGTTCGTCACATCTCATCTACCTCCCGGTTTTAATGAATACGATTTTGTGCCAGAGTCCTTCGATAGGGACAAGACAATTGCACTGATCATGAACTCCTCTGGATCTACTGGTCTGCCTAAAGGTGTCGCTCTGCCTCATAGAACTGCCTGCGTGAGATT | Sigma | Cat#EHUFLUC |
| EHD2 Fw (HindIII):  CGCAAAGCTTCTATGTTCAGCTGGCTGAAGCGG | Integrated DNA Technologies | N/A |
| EHD2 Rev (ECORI):  ATACGAATTCTCTCGGCGGAGCCCTTGTGGCG | Integrated DNA Technologies | N/A |
| Software and Algorithms | | |
| ImageJ | N/A | https://Imagej.nih.gov/ij |
| MaxQuant (version 1.6.7.0) | N/A | https://www.maxquant.org/ |
| Uniprot | N/A | https://www.uniprot.org/ |
| ProTIGY | Broad Institute, Proteomics Platform | https://github.com/broadinstitute/protigy |
| Cytoscape | N/A | https://cytoscape.org |
| BioGRID | N/A | https://thebiogrid.org |
| STRING | N/A | https://string-db.org |
| Inkscape | N/A | https://inkscape.org/ |
| R (version 4.0.3) | N/A | https://www.r-project.org/ |
